# Generation and Validation of Versatile Inducible CRISPRi Embryonic Stem Cell and Mouse Models

**DOI:** 10.1101/2020.04.28.065854

**Authors:** Rui Li, Xianyou Xia, Xing Wang, Xiaoyu Sun, Zhongye Dai, Dawei Huo, Huimin Zheng, Haiqing Xiong, Aibin He, Xudong Wu

## Abstract

CRISPR-Cas9 has been widely used far beyond genome editing. Fusions of deactivated Cas9 (dCas9) to transcription effectors enable interrogation of the epigenome and controlling of gene expression. However the large transgene size of dCas9-fusion hinders its applications especially in somatic tissues. Here, we develop a robust CRISPR interference (CRISPRi) system by transgenic expression of doxycycline (Dox) inducible dCas9-KRAB in mouse embryonic stem cells (iKRAB ESC). After introduction of specific gRNAs, the induced dCas9-KRAB efficiently maintains gene inactivation, though it exerts modest effects on active gene expression. Proper timing of Dox addition during cell differentiation or reprogramming allows us to study or screen spatiotemporally activated promoters or enhancers and thereby the gene functions. Furthermore, taking the ESC for blastocyst injection, we generate an iKRAB knockin (KI) mouse model that enables shut-down of gene expression and loss-of-function studies *ex vivo* and *in vivo* by a simple transduction of gRNAs. Thus, our inducible CRISPRi ESC line and KI mouse provide versatile and convenient platforms for functional interrogation and high-throughput screens of specific genes and potential regulatory elements in the setting of development or diseases.

## Introduction

Clustered regularly interspaced short palindromic repeat (CRISPR) and Cas (CRISPR- associated) proteins were originally found in bacteria and archaea to defend against viruses and plasmids by using CRISPR RNAs to guide the silencing of invading nucleic acids. Rapidly, this system has been simulated in other species by introducing the endonuclease Cas9 and single guide RNAs (sgRNAs) to cleave and edit specific DNA sequences. Ever since, the (CRISPR)/Cas9 system has been widely used as a powerful tool for genome editing [1-4]. Meanwhile, the RNA-guided epigenome editing technologies based on endonuclease deactivated Cas9 (dCas9) with two point mutations (D10A, H841A) have been developed. By fusion of dCas9 with transcription activator or repressor, the CRISPR activation (CRISPRa) or CRISPR interference (CRISPRi) allows researchers to control the level of endogenous gene expression [5-9].

Conventional genome editing by CRISPR/Cas9 technology may result in divergent indels and generate differential genotypes. In comparison, CRISPRi techniques block gene transcription by introducing transcription repressors at a defined genomic locus, leaving DNA sequence intact. Krüppel-associated box (KRAB) domain are the most commonly used repressors [6, 7, 10]. Epigenetic studies have demonstrated that KRAB containing zinc-finger proteins (KRAB-ZFPs) facilitate silencing by recruiting the KRAB associated protein KAP1, and in turn other epigenetic repressors such as SETDB1, EHMT2/G9A, LSD1 and NURD complex. Thus the dCas9-KRAB fusion protein creates an inactive chromatin environment by removing active chromatin mark like histone H3-acetylation and establishing heterochromatin-like chromatin mark H3 lysine 9 trimethylation (H3K9me3) [11-14].

Current CRISPRi systems are usually generated by ectopic expression of dCas9-KRAB and sgRNAs via viral transduction. However the expression of bacterial Cas9 could elicit host responses, aberrant cellular functions or even toxicity in mammalian tissues [15, 16]. Considering these potentially detrimental effects, lasting expression of Cas9 or dCas9 proteins is not preferred. Besides, controllable genetic manipulation is crucial for most of the biological studies. Hence the inducible expression of dCas9-KRAB serves as a better choice. Here we generated a transgenic mouse embryonic stem cell (ESC) line with doxycycline (Dox) inducible and reversible expression of dCas9-KRAB (iKRAB ESC). With this line, a simple transduction of sgRNAs enables us to do any locus-specific loss-of-function (LOF) studies in ESC or differentiated cells by proper timing of Dox addition or withdrawal. Having tested multiple gRNAs targeting promoters or enhancers, we found that our iKRAB system could efficiently maintain gene inactivation and control cell fate transition. And high-throughput screens could be performed in iKRAB ESC-derived cells. To facilitate broader applications, we took advantage of the ESC line and generated knockin mice, which allowed inducible LOF studies *ex vivo* and *in vivo*. These systems especially the animal models empower us for functional studies and potentially high-throughput screens of specific genes or *cis*-regulatory elements.

## Results

### Generation and characterization of iKRAB ESC line

To generate a robust and inducible CRISPRi system, we took advantage of an engineered inducible cassette exchange (ICE) system of mouse ESCs (A2Loxcre) with rtTA inserted into the *Rosa26* locus and TRE-LoxP-Cre-LoxP-Δneo integrated at the housekeeping gene *Hprt* [17]. Transgenes integrated at the *Hprt* locus remain transcriptionally active in differentiated cell types as well as in ESC. First we constructed a dCas9-KRAB fragment onto the p2Lox- FLAG vector which contains the LoxP sites [18]. Then we pre-treated A2Loxcre cells with Dox for 16 hour (hrs) so that *Cre* is expressed and the cells are competent for recombination. Upon transfection with the p2Lox-FLAG-dCas9-KRAB construct, homologous recombination was initiated at the LoxP locus, and genomic fragments coming from plasmids were integrated into the downstream of the Tetracyclin Response Element (TRE) promoter. At the same time, the Δneo gene acquired a *PGK* promoter and a start codon and enabled us to select for the precise integration with G418 (Fig 1A). After around 10 days’ selection, the resistant clones were picked and characterized by genotyping PCR analysis. Two positive clones showed that the FLAG-dCas9-KRAB expressing sequence was precisely integrated downstream of TRE (S1A Fig). One of the clones was expanded for further analysis.

**Fig 1.**
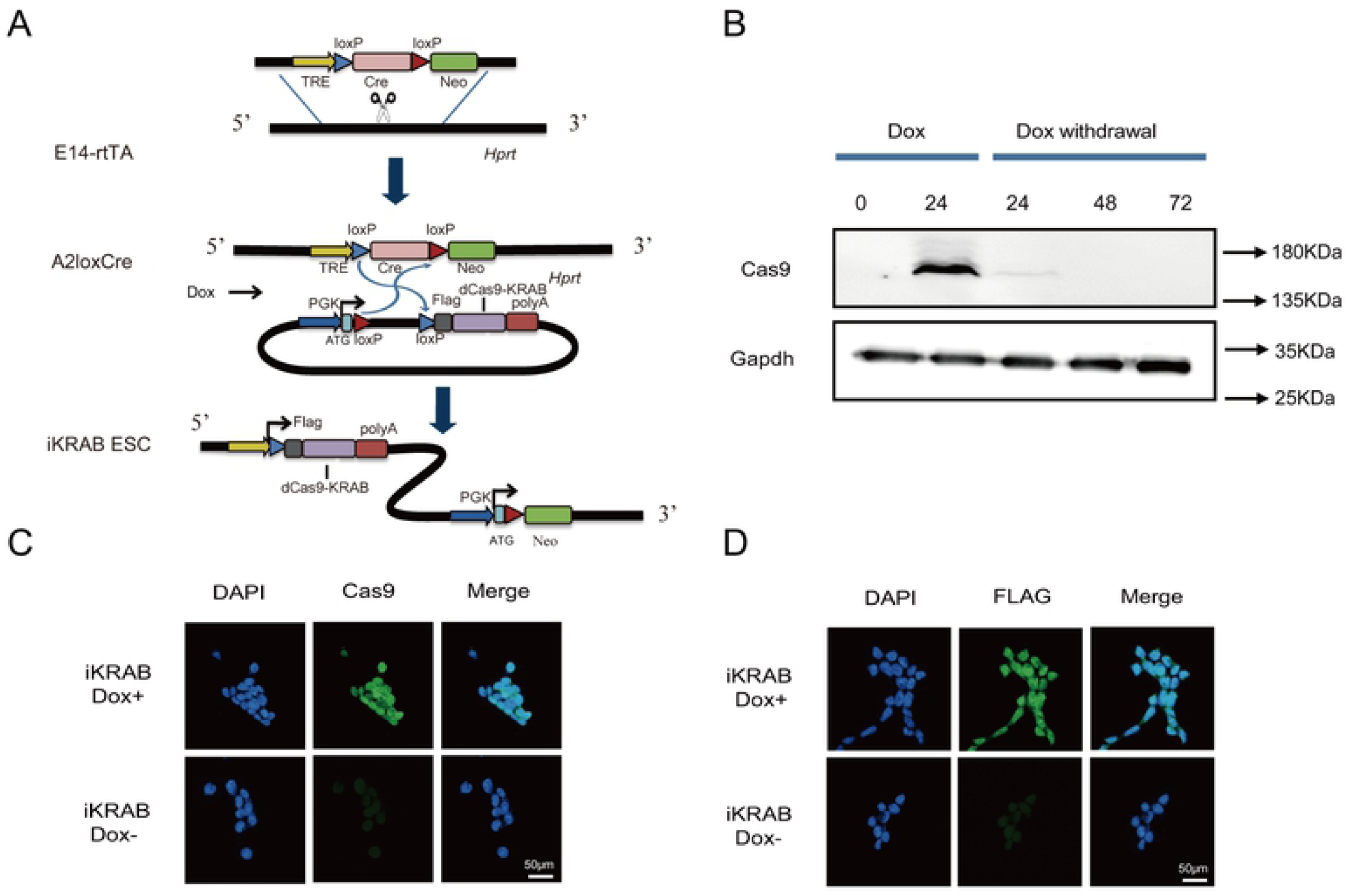
Generation of the iKRAB ESC line. (A) Schematic diagram shows the strategy of inducible cassette exchange to generate the iKRAB ESC line. FLAG-dCas9-KRAB was integrated into the downstream of the TRE element through homologous recombination. Dox- controlled reverse transcriptional activator (rtTA) drives expression of fusion protein of FLAG-dCas9-KRAB. (B) Western blot analysis showing the inducible and reversible expression of FLAG-dCas9-KRAB protein at different time points after Dox addition or withdrawal. β-actin served as a loading control. (C-D) Immunofluorescence staining of Cas9 and FLAG in iKRAB ESC cultured with or without Dox. The scale bar represents 50 μm.

As examined by Western Blot assay with the Cas9 antibody, the clone did not express any detectable dCas9-KRAB protein when cultured without Dox, indicating no leaky expression. Upon addition of titrated concentration of Dox, dCas9-KRAB expression was robustly induced at 1 μg/ml after 24 hrs (S1B Fig and Fig 1B). Hereafter we used Dox at 1 μg/ml for most of the experiments unless otherwise stated. Interestingly, the protein expression was gradually decreased to undetectable level 48 hrs after removing Dox (Fig 1B). Hence we named this clone iKRAB ESC. Meanwhile, we did Immunofluorescence (IF) analysis of the clone with Cas9 and FLAG antibodies. As shown in Fig 1C & 1D, the fusion protein was homogenously expressed after Dox treatment. These data showed that dCas9- KRAB expression could be precisely and reversibly controlled by the addition and withdrawal of Dox, which would allow controllable gene knockdown upon introduction of gRNAs.

### iKRAB efficiently maintains rather than induces gene silencing

Many previous studies have demonstrated that dCas9-KRAB can achieve efficient knockdown of gene transcription, especially when tethered near the transcription start site (TSS) by sgRNAs [6, 8, 9, 19]. To test the efficiency of the iKRAB ESC line, we first designed six specific sgRNAs targeting near the TSS of *Oct4*. As Oct4 is one of the best known pluripotency factors and required for ESC self-renewal, its depletion is expected to result in a clear loss of pluripotent cell morphology. However unexpectedly, quantitative reverse transcription PCR (RT-qPCR) analysis showed that none of the sgRNAs downregulated the mRNA levels of Oct4 more than 50% after Dox induction, no matter targeting upstream or downstream of the TSS (Fig 2A). The efficiency was not improved even we designed more sgRNAs according to the FANTOM5/CAGE-defined TSS [19] (data not shown). And IF analysis of cells transduced by Oct4#sgRNA4, the most effective one in our test, clearly showed that Oct4 expression was homogenously downregulated after Dox treatment (S2A Fig). Thus the poor efficiency was not due to heterogeneous expression of sgRNAs or inadequate transduction rate. Similar CRISPRi inefficiency was also observed for other actively expressed genes in ESCs such as *Bap1, Kdm2b*, etc (S2B Fig and data not shown). Therefore our data argue against the high CRISPRi efficiency by iKRAB on actively transcribed genes.

**Fig 2.**
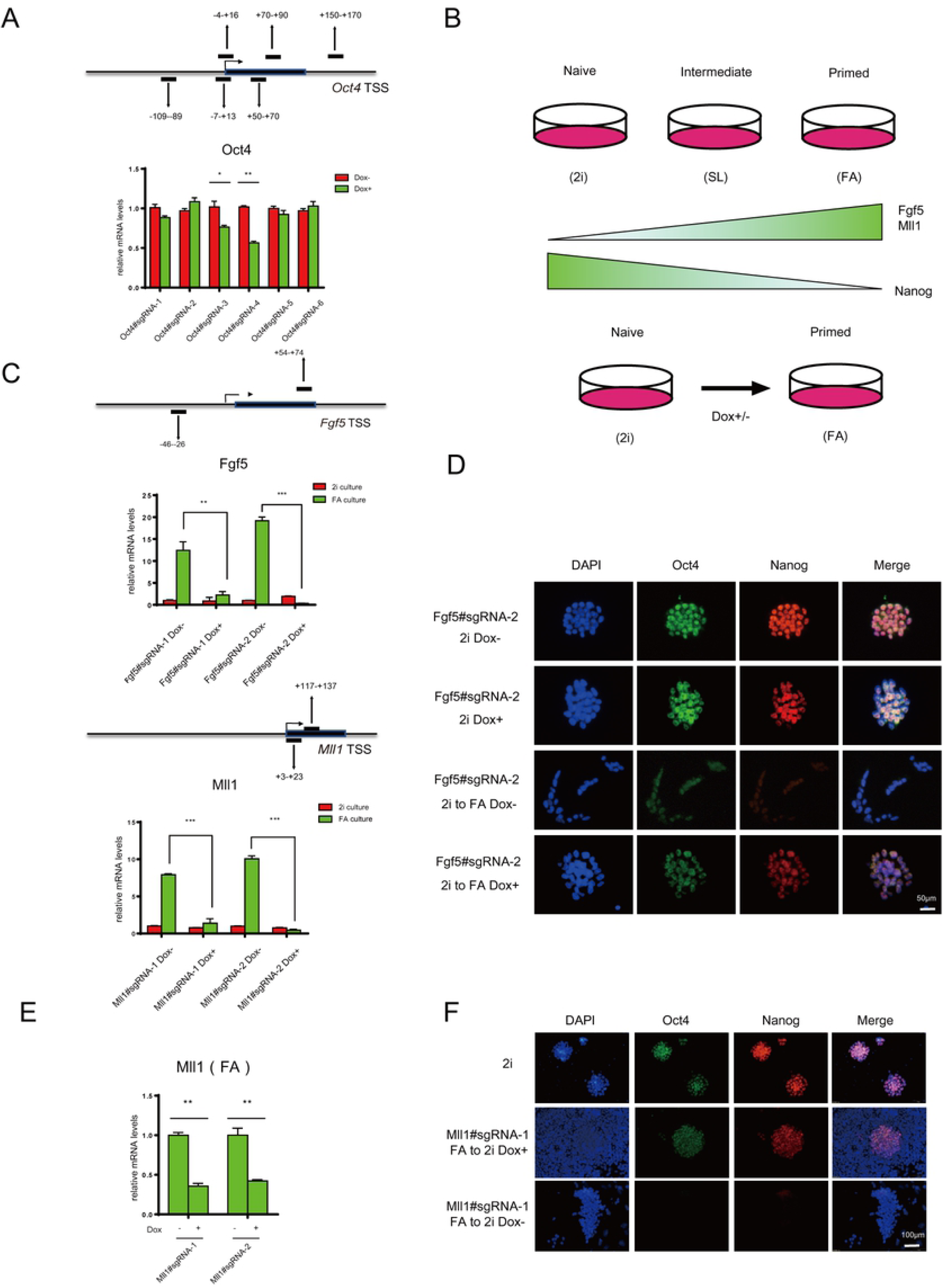
Induced dCas9-KRAB at promoters is sufficient to maintain rather than induce gene inactivation. (A) RT-qPCR analysis of stable iKRAB ESCs containing sgRNA against *Oct4* showed less than 50% knockdown efficiency after 2 days of Dox induction. The binding location of each sgRNA is indicated relative to the TSS of *Oct4* locus. (B) Schematic diagram shows the dynamics of different states and marker gene expression of pluripotent stem cells cultured in different conditions. (C) RT-qPCR analysis of Fgf5 and Mll1 mRNA levels of iKRAB cells containing designated sgRNAs (with or without Dox) upon culture condition switch from 2i to FA. The binding location of each sgRNA is indicated relative to the TSS of the *Fgf5* or *Mll1* locus. (D) Immunofluorescence staining of Oct4 and Nanog in iKRAB cells containing Klf5#sgRNA-2 treated with or without Dox in 2i condition or 2i switch to FA condition. The scale bar represents 100 μm. (E) RT-qPCR analysis of Mll1 mRNA levels of iKRAB cells containing the same sgRNAs as (C) (with or without Dox) in FA condition. (F) Immunofluorescence staining of Oct4 and Nanog in iKRAB cells containing Mll1#sgRNA-1 treated with or without Dox for 4 days in FA condition, followed by switch back to 2i condition. ESC constantly cultured in 2i condition was used as a control. The scale bar represents 100 μm. Data in (A, C, E) are represented as the mean ± SD of replicates (n=3). * p<0.05, ** p<0.01, ***p<0.001; and two-tailed unpaired *t* test.

A main advantage of an inducible system is to precisely control gene expression via the timing of Dox addition. Since the iKRAB was not sufficient to induce gene inactivation, we followed to test whether it would maintain gene inactivation in dynamic settings. We took advantage of the switch between naïve and primed pluripotent states of iKRAB ESC. ESCs cultured in serum-free medium containing GSK3 inhibitor and MEK inhibitor (2i) and LIF (hereafter 2i) supports naïve pluripotency that mimics the inner cell mass (ICM) of the blastocyst. Upon switch to serum-containing medium with LIF (hereafter SL) or Fgf2 and activin (hereafter FA), ESCs will exit from naïve pluripotency and switch to primed pluripotency. Some factors such as the epiblast maker Fgf5 and the histone methyltransferase Mll1 are switched on during the transition, accompanied with loss of Nanog expression [20-22] (Fig 2B). We transduced iKRAB ESCs with lentivirus expressing sgRNAs targeting the TSS of *Fgf5 or Mll1*. In the absence of Dox, *Fgf5* or *Mll1* expression was strongly induced upon the switch of the culture condition from 2i to SL or FA. Though Dox addition in 2i medium exerted minor effects on *Fgf5* or *Mll1* expression, it abolished the induction by medium switch (>85%) (Fig 2C). As shown by the cell morphology and immunostaining of Oct4 and Nanog, the Dox-treated cells maintained naïve state (round colonies, Oct4 and Nanog positive) in response to FA signals, indicating that Fgf5 inactivation did hinder the exit of naïve pluripotency (Fig 2D).

Notably, Chromatin Immunoprecipitation (ChIP)-qPCR analysis in Oct4#sgRNA4 and Fgf5#sgRNA1 transduced ESCs showed that the efficiency of tethering dCas9-KRAB at the designed locus in Dox-treated cells was comparable (S3 Fig). These data suggested that the divergent effects on active *versus* inactive genes were not due to the differential genome binding of dCas9-KRAB protein. Moreover, when we initiated Dox treatment at the primed state with *Mll1* highly expressed (FA culture), the same sgRNAs targeting the TSS of *Mll1* worked less efficiently (∼60% downregulation) (Fig 2E). Nonetheless, upon being switched back to 2i medium, some of the Mll1 knockdown cells were successfully reprogrammed from primed state to naïve state as previously reported [22] (Fig 2F). This finding also prompts us for further CRISPRi screening to identify more chromatin regulators whose suppression may contribute to the reprogramming. Together these data demonstrate that the iKRAB system is highly efficient to block gene activation though insufficient to induce silencing of active genes.

Nevertheless, highly efficient gene silencing is not always preferred, especially to genes critical for cell survival. For example, *Bap1* deletion in ESCs triggers apoptosis [23] and hence CRISPRi may suffice its LOF studies. Moreover, different levels of gene downregulation may help get insight into gene dosage effect. Interestingly, we observed that the ratio of Nanog-positive cells was even increased in the above SL-cultured Oct4-CRISPRi ESCs, suggesting a robust pluripotent state (S2A Fig). This is consistent with a previous unexpected finding that the self-renewal efficiency in Oct4+/- ESCs is even enhanced compared with the wild type (WT) [24]. Accordingly, the iKRAB system may provide a valuable means to precisely control the dosage as well as the timing of target gene expression.

### iKRAB maintains inactive enhancers

Epigenome editing is supposed to modulate activities of any potential *cis*-acting regulatory elements. Enhancers are a vital regulatory element for tissue or development stage-specific gene expression through interaction with promoters. Thus we tested how the iKRAB system works at enhancers in ESC and derived cells. An attractive model is the dynamic reorganization of enhancers between the two states of pluripotent stem cells [25, 26]. For example, *Oct4* expression is controlled by different enhancers, distal enhancer (DE) in naïve state while proximal enhancer (PE) in primed state, though *Oct4* is expressed in both naïve and primed pluripotent cells [25, 27]. When two sgRNAs targeting *Oct4* PE were respectively introduced into the 2i cultured iKRAB ESCs, Dox treatment induced no effects or even slight upregulation of Oct4 expression, as shown by RT-qPCR analysis (Fig 3A and S4 Fig). Upon the culture condition switching from 2i to either SL or FA without Dox, Oct4 expression levels were increased in SL condition, an intermediate state with simultaneous activation of two enhancers. However after Dox treatment, Oct4 activation was successfully suppressed in SL condition, and was almost abrogated in FA condition (Fig 3B). IF analysis further confirmed that Oct4 expression was impeded in FA condition at the presence of Dox, though it was unaffected in 2i condition. Moreover, we found that Oct4 expression was restored 4 days after withdrawal of Dox, indicating CRISPRi effect as well as the expression of dCas9- KRAB fusion protein was reversible (Fig 3C). ChIP-qPCR analysis of 2i and FA-cultured cells showed that H3K27me3 and H3K9me3 levels at *Oct4* PE were decreased, together with increased H3K4me1 and H3K27ac levels upon switch from 2i to FA condition. However when culture condition was switched at the presence of Dox, H3K27me3 and H3K9me3 levels at *Oct4* PE were maintained or even increased and the increase of H3K4me1 and H3K27ac levels was hindered (S5 Fig). These data clearly indicated that *Oct4* PE was blocked at inactive state by induced dCas9-KRAB.

**Fig 3.**
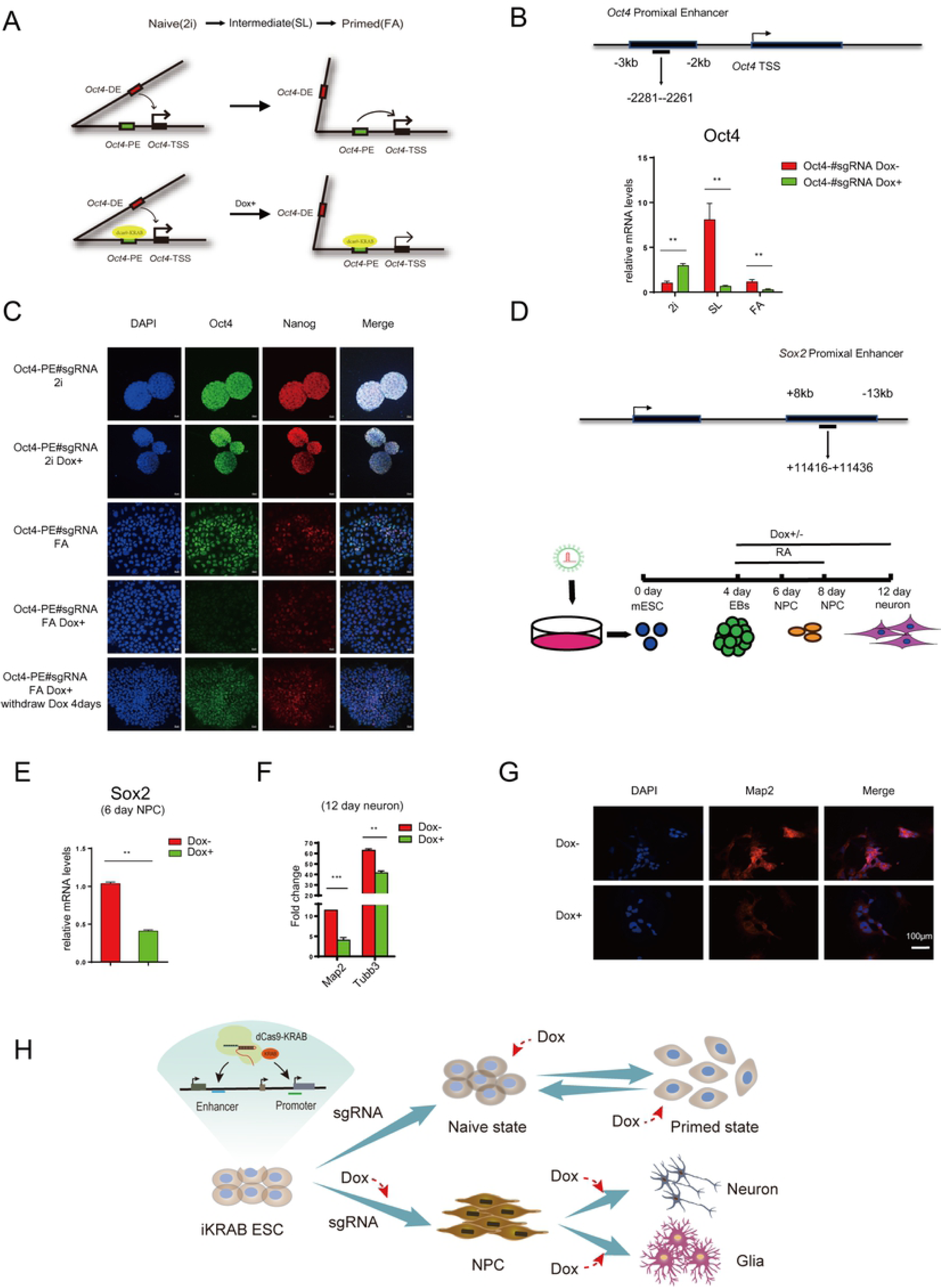
Induced dCas9-KRAB at enhancers is sufficient to maintain gene inactivation in response to activation signals. (A) Schematic diagram shows the proximal enhancer of *Oct4* is activated upon culture condition switch from 2i to FA. Dox-induced dCas9-KRAB is tethered to the proximal enhancer of *Oct4* in 2i condition and Oct4 expression is to be tested in SL and FA condition. (B) RT-qPCR analysis of Oct4 mRNA levels of iKRAB cells containing designated sgRNAs (with or without Dox) upon culture condition switch from 2i to SL or FA. The binding location of each sgRNA is indicated relative to the proximal enhancer of *Oct4* locus. (C) Immunofluorescence staining of Oct4 and Nanog in designated conditions. The scale bar represents 20 μm. (D) Schematic diagram shows iKRAB cells containing designated sgRNAs the proximal enhancer of *Sox2* are differentiated to NPC and neuron. Dox was added before NPC stage together with RA. (E) RT-qPCR analysis of Sox2 mRNA levels in NPC (6 days) from the group with or without treatment. (F) RT-qPCR analysis of two neuron marker genes in neuron (12 days) from the group with or without treatment. (G) Immunofluorescence staining of Map2 in neuron (12 days) from the group with or without treatment. The scale bar represents 50 μm. (H) A model for the temporal control of enhancer or promoter by the timing of Dox addition during ESC differentiation. Data in (B, E, F) are represented as the mean ± SD of replicates (n=3) (** p<0.01, ***P<0.001 and two-tailed unpaired *t* test).

To further test the eligibility of iKRAB ESC for the dissection of specific enhancers during differentiation, we established a neural differentiation model to observe the dynamic control of *Sox2* enhancers and its downstream effects. Sox2 is highly expressed in Neural Progenitor Cells (NPCs) as well as in ESCs, however its expression is activated by distal enhancer (DE) in ESCs while likely by proximal enhancer (PE) in NPCs [28, 29]. We transduced the iKRAB ESCs with lentivirus expressing a specific sgRNA targeting *Sox2* PE before proceeding with embryoid body (EB) differentiation. When RA-induced NPC differentiation was initiated, cells were cultured with or without Dox (Fig 3D). RT-qPCR analysis in 6 days differentiated NPCs showed that Sox2 mRNA levels were significantly decreased in the Dox-treated group compared with the mock control, indicating that the switch-on of *Sox2* PE was blocked (Fig 3E). Consistent with the activation of *Sox2* PE in NPCs, the interaction between *Sox2* PE and TSS was increased after RA treatment, as shown by the 3C-PCR analysis. And this interaction failed to be established in the Dox-treated cells (S6 Fig), suggesting that induced dCas9-KRAB suppressed *Sox2* PE activation and chromatin looping. Then we continued the differentiation of NPCs to neuron for each group and observed how Sox2 inactivation affected sequential neurogenesis. RT-PCR analysis showed that the expression levels of two neuron marker genes *Map2* and *Tubb3* were significantly lower in the Dox-treated group than the control group (Fig 3F). The inadequate Map2 expression in the Dox-treated group was further confirmed by IF analysis, indicating differentiation defects (Fig 3G). Collectively, a simple introduction of sgRNAs in iKRAB ESC can efficiently guide temporally induced dCas9-KRAB to block the activation of lineage-specific gene promoters or enhancers and thereby affect cell fate transitions (Fig 3H).

### iKRAB ESC for high-throughput screening

As ESCs have the potential to differentiate into all cell types of the organism, the iKRAB ESC is supposed to be optimal for the identification of specific promoters or enhancers and characterization of associated genes at defined contexts. And Hence we developed a CRISPRi screen to identify specific chromatin regulators whose inhibition would alleviate the toxicity of sodium channel blockers in the derived neural cells. First we created a sgRNA library (containing 5,115 sgRNAs with 5,096 specific sgRNAs and 19 non-targeting negative control sgRNAs) that targeted the TSSs (including different transcripts) of known or potential epigenetic regulators and RNA binding proteins coding genes (total number=857). Then we transduced the sgRNA library into the iKRAB ESC, induced CRISPRi activity before differentiation to NPC stage while treated the cells with NaV1.8 channel blocker A803467. The Dox-free group of NPCs almost completely died after 4 days of A803467 treatment. The survived cells in Dox-treated group were harvested for deep sequencing (Fig 4A).

**Fig 4.**
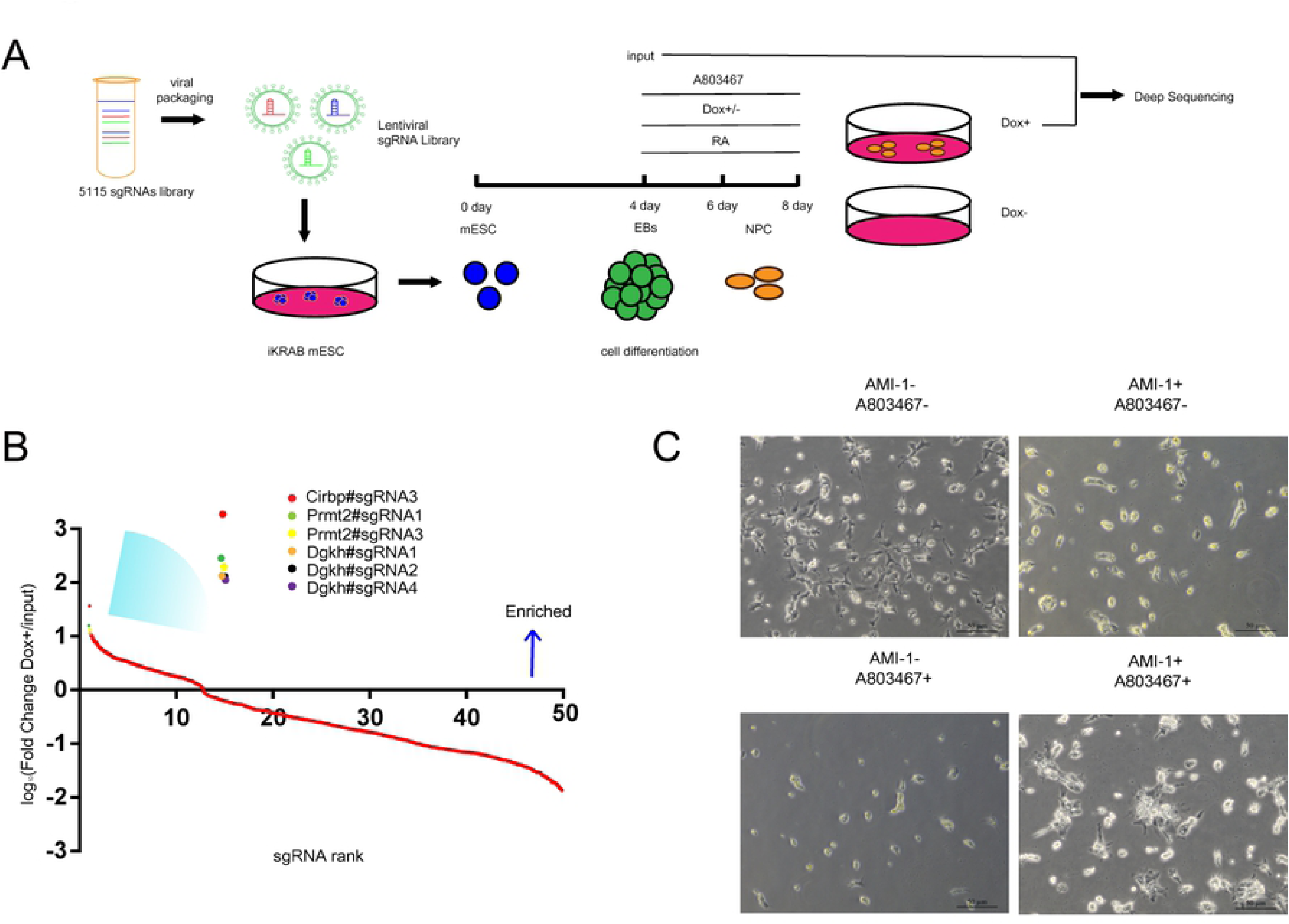
iKRAB ESC for screening. (A) Schematic representation of CRISPRi screens in the iKRAB ESC-derived neural cells to evaluate chromatin regulators whose depletion would resist the toxicity of sodium channel blockers A803467. (B) A map of the contribution of the top 414 sgRNAs enriched in A803467-resistant cells. PRMT2 is among the top hits. (C) Validation the neuroprotective effects of PRMT inhibitor AMI-1. The scale bars represent 50 μm

Compared with the input, 41 sgRNAs for 19 genes were significantly enriched in the survived cells (log10(Fold change Dox+/input)>1, S1 Table). *Cirbp, Prmt2* and *Dgkh* were among the top hits (Fig 4B). PRMT2 belongs to type I Protein Arginine Methyltransferases and there were inhibitors available against its activity, such as AMI-1. So we examined whether AMI-1 would alleviate the toxicity of A803467 in NPCs. As shown in Fig 4C, AMI- 1 significantly promoted NPC survival and partly maintained the cell morphology at the presence of A803467. Accordingly, the CRISPRi screening identified toxicity resistance gene and provided possible solutions for neuroprotection. Thus the iKRAB ESC may serve as a versatile model for LOF screens of functional genes or regulatory elements in a wide range of settings.

### Generation of an iKRAB KI mouse model for inducible gene silencing *ex vivo* and *in vivo*

Though dCas9-KRAB has been broadly applied in a few cell lines [8, 9, 30, 31], direct applications in somatic tissues remain challenging, mainly owing to its large transgene size. A robust *in vivo* CRISPRi system is urgently needed. Since we have confirmed CRISPRi effects of the iKRAB ESC, we performed blastocyst injection. After successfully obtaining mouse chimeras and screening of germline transmitted offsprings, chimeric founder mice were crossed to generate iKRAB heterozygous or homozygous KI mice, verified by genotyping PCR (Fig 5A and 5B). The iKRAB KI mice were fertile, presented no morphological abnormalities, and were able to breed to homozygosity. Western Blot assay of protein lysates from the mouse tails showed that no expression of dCas9-KRAB was observed in the KI mice until being fed with Dox-containing water (Fig 5C). Then we tested inducible CRISPRi effect *ex vivo* and *in vivo*.

**Fig 5.**
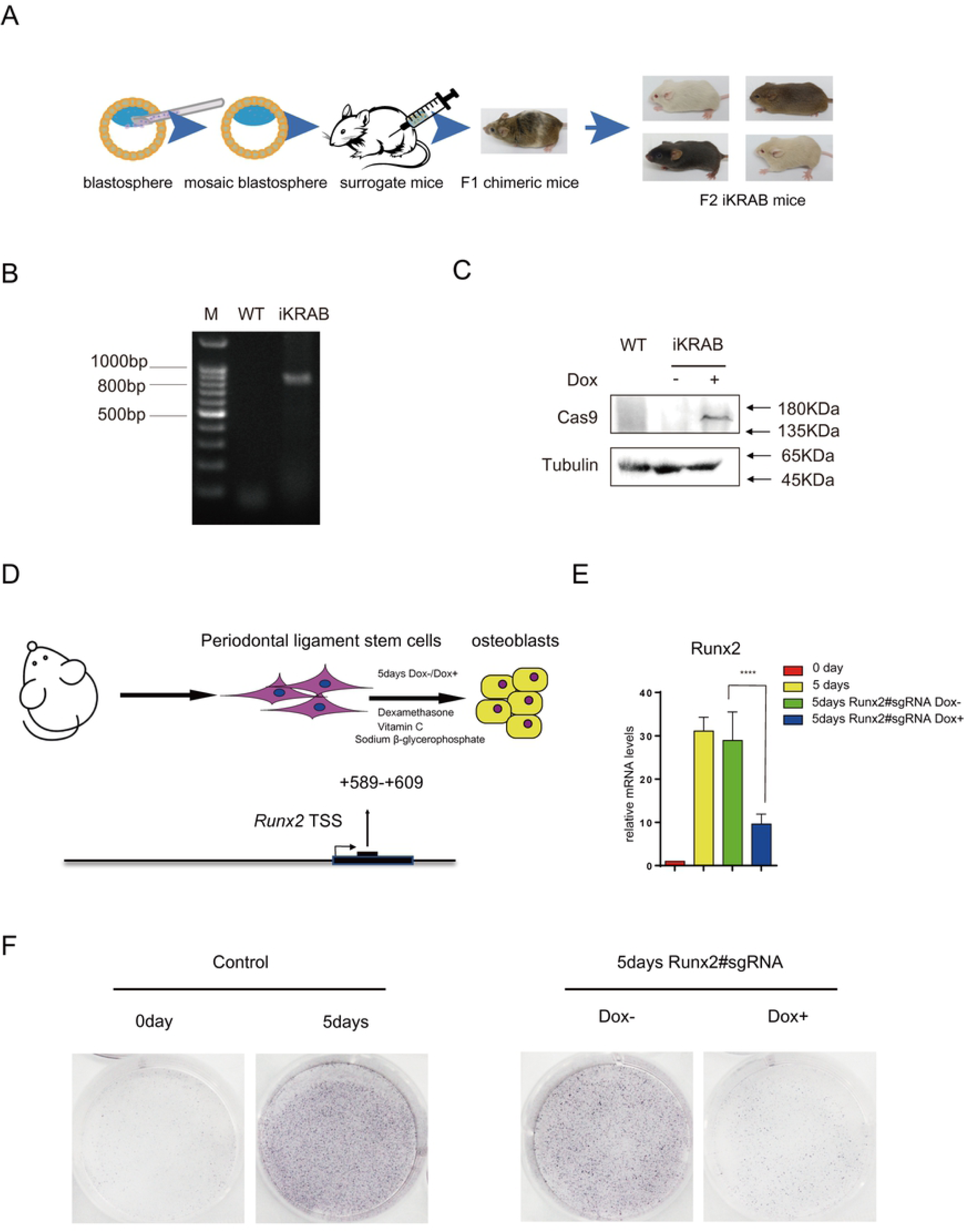
Characterization of the iKRAB KI mouse and *ex vivo* effect. (A) Schematic diagram shows the generation of iKRAB KI mice. (B) Genotyping PCR analysis of WT and iKRAB KI mice. (C) Western blot analysis of dCas9-KRAB expression in WT and iKRAB mice. β-tubulin served as a loading control (D) Schematic diagram shows that Periodontal ligament stem cells from the iKRAB KI mice was introduced with sgRNA against the TSS of *Runx2*, followed by differentiation into osteoblasts. (E) RT-qPCR analysis showing Runx2 mRNA levels in the designated groups. Data are represented as the mean ± SD of replicates (n= 3) (**** p<0.0001; and two-tailed unpaired *t* test). (F) ALP staining of cells from the designated groups.

To test the CRISPRi effect *ex vivo*, we took advantage of a previously established osteoblast differentiation model of mesenchymal stem cells (MSC) [32]. Periodontal Ligament Stem Cells (PDLSCs) were harvested from 8 weeks old iKRAB KI mice. After a short expansion in MSC medium, the cells were switched to differentiation medium (Fig 5D). RT-qPCR analysis showed that Runx2 expression was activated after 5 days’ differentiation (Fig 5E). Meanwhile, alkaline phosphatase (ALP) activity, an early marker for osteoblast differentiation, was strongly induced (Fig 5F). Then we transduced PDLSCs with lentivirus expressing specific sgRNA targeting the TSS of *Runx2*, encoding a key transcription factor driving osteoblast differentiation. The transduced cells were then switched to differentiation medium with or without Dox. We found that Dox treatment significantly inhibited the differentiation medium-induced Runx2 activation (Fig 5E) and ALP activity (Fig 5F). These data illustrate that primary cells isolated from the iKRAB KI mice respond well to Dox induction.

To directly test the CRISPRi effect *in vivo*, we designed three sgRNAs, which respectively target the specific enhancer and the first exon of *Mitochondrial transcription factor A* (*TFAM*), whose depletion results in muscle atrophy [33]. To achieve high knockdown efficiency *in vivo*, we cloned multiplex gRNAs by linking the three sgRNAs linearly into an adeno-associated viral (AAV) vector simultaneously expressing GFP. The construct was used for AAV packaging and purification. Then the high titer virus was injected into *Tibialis anterior* muscle of 6 weeks old iKRAB KI mice. Dox-containing water was fed to the mice two weeks after injection. We chose to test in muscles mainly considering of technical feasibility and local virus concentration. Mice were sacrificed after 1 month of Dox induction and tissues from *Tibialis anterior* muscle were isolated for analysis (Fig 6A). Successful transduction was confirmed by the observation of GFP. And Tfam expression levels were significantly lower in GFP+ cells than in the GFP- cells as shown by RT-qPCR analysis of the sorted cells (Fig 6B). IF analysis of GFP and laminin expression at the myofiber membrane demonstrated that the diameter of muscle fibers was significantly smaller in the AAV infected muscle fibers (GFP+) compared with the un-infected ones (GFP-) (Fig 6B). The cross-sectional areas of the GFP+ with GFP- myofibers from six mice were calculated. As shown in Fig 6C, the muscle fibers with downregulated Tfam expression were much smaller than the control group, indicative of muscle atrophy. Taken together, the iKRAB KI mice provide a versatile model for ex *vivo* and *in vivo* LOF studies.

**Fig 6.**
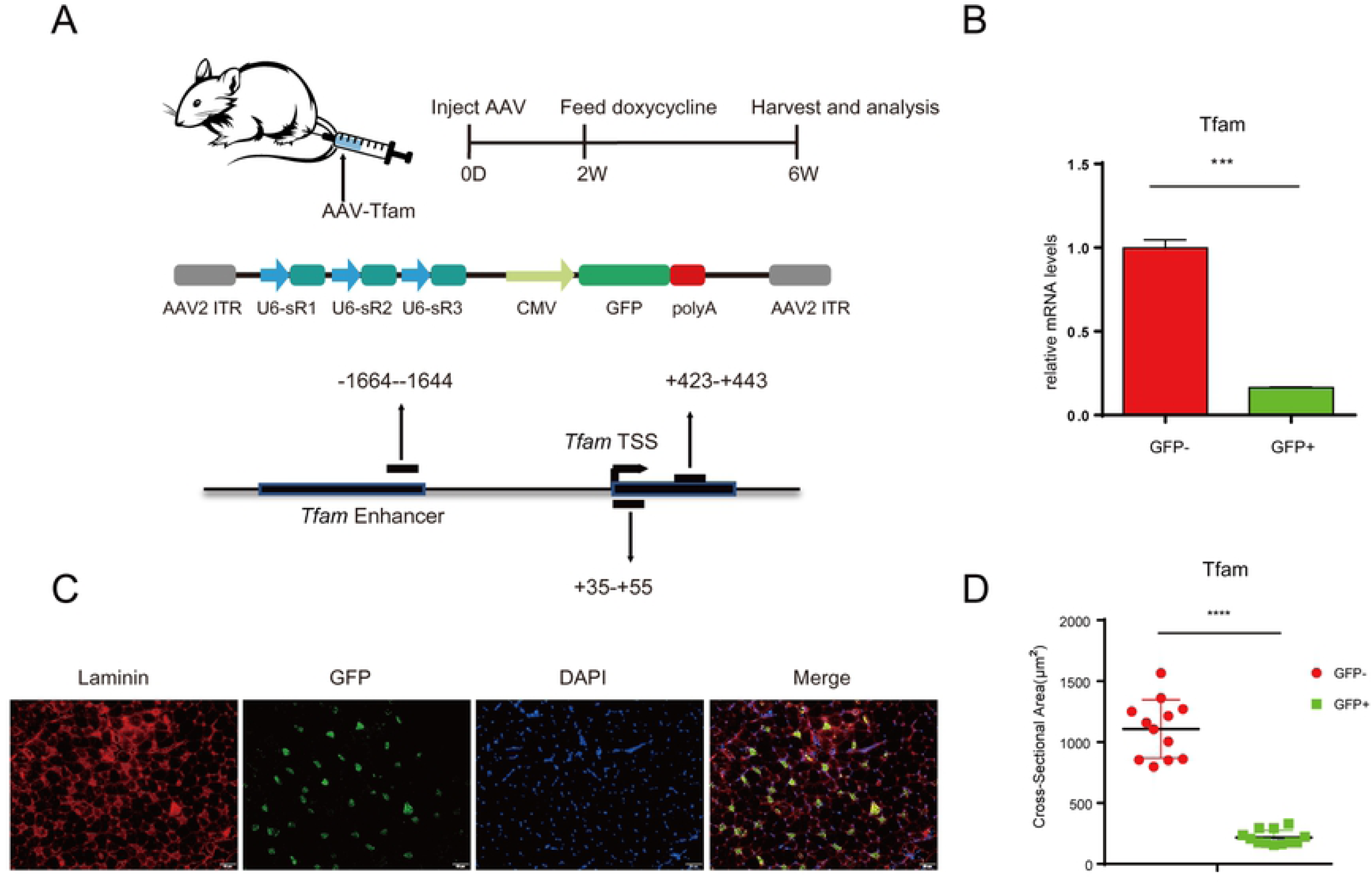
*In vivo* inducible CRISPRi effect of the iKRAB KI mouse. (A) Schematic diagram shows that AAV expressing multiplex gRNAs against Tfam locus was injected into *Tibialis anterior* muscle of the iKRAB KI mice, followed by 1 month of Dox induction and the subsequent analysis. (B) RT-qPCR analysis showing Tfam mRNA levels in the designated groups. Data are represented as the mean ± SD of replicates (n= 3) (*** p<0.001; and two- tailed unpaired *t* test) (C) Immuno?uorescence analysis of Laminin of *Tibialis anterior* muscle tissue around the injection sites. GFPs mark the infected muscle fibers. The scale bar represents 50 μm. (D) Quantification of the cross-sectional area of *Tibialis anterior* muscle fibers (n=6 mice). The data are presented as means ± SD (****p < 0.0001; and two-tailed unpaired *t* test).

## Discussion

In this study, we generate an inducible CRISPRi mouse ESC line and KI mouse, inducibly and reversibly expressing dCas9-KRAB protein. With either the cell line or animal model, a simple transduction of gRNAs enables us to do any controllable LOF studies.

CRISPRi techniques have been shown to repress gene transcription to different extent in a variety of cell models [34]. Here we argued that induced dCas9-KRAB displayed limited or no effects on active promoters or enhancers. And it was not due to un-optimized gRNA designing because even the same sgRNA achieved differenial repressive activities at distinct cellular states (Fig 2). Actually this issue was also recently raised by several other studies [35-37]. For example, the sgRNA targeting the *MYC* promoter that led to downregulated MYC expression 6.2 fold in HEK293T cells showed very modest or no repressive activity in cancer cell lines with high MYC expression [36]. This performance is consistent with the transcriptional repression mechanism that KRAB-ZFPs facilitate heterochromatin formation and spreading at inactive chromatin regions [11, 13, 14, 38]. Though dCas9-KRAB protein is tethered to chromatin locally by sgRNAs, it is not sufficient to counteract the active chromatin environment for the propagation of heterochromatin-like features. And we did find that multiple gRNAs targeting a wide-range of regions at the same gene (e.g., Tfam) mediated sufficient transcriptional repression. Considering of the increased risk of off-target of multiplex gRNAs, other optimization strategies need to be tested to improve CRISPRi effect on active genes in the future. Recently KRAB combination with MeCP2 or LSD1 or alternative repressors like SIN3-interacting domain (SID) have been reported to achieve superior efficiency [35, 37, 39] and are worth further testing at active promoters or enhancers in different cell types.

Despite of insufficiency to induce silencing, we showed that dCas9-KRAB pre-set at inactive promoters or enhancers was sufficient to maintain or foster silencing. Taking cell differentiation or reprogramming models, we demonstrated that Dox-induced locus-specific perturbation in iKRAB cells is competent to restrict gene activation and thereby affect cell fate transitions. Moreover, though we only test gRNAs targeting a single gene in our study, we believe that simultaneous introduction of sgRNAs targeting multiple genes would work as well.

In addition to functional studies of individual genes or *cis*-regulatory elements, genome-scale CRISPRi screens have been widely applied to identify genes or noncoding RNAs that control diversity of cellular processes [8, 31, 35, 40, 41]. Similarly, we can take iKRAB ESC or its derivatives for CRISPRi screens to identify new genes that regulate stem cell self-renewal and differentiation, to screen potential barriers against reprogramming, or to map any key *cis*-regulatory elements especially enhancers for cell fate decisions (Fig 3H). For this purpose, the iKRAB KI mouse will provide a convenient platform for broader applications. Taking adult stem cells or differentiated cells (e.g., fibroblast) from the iKRAB KI mice for *ex vivo* functional studies or screens will avoid the problems of inadequate ESC differentiation, which will consequently contribute to the improvement of differentiation or reprogramming efficiency for regenerative medicine in the long term. For cellular processes like hematopoiesis, a more physiologically relevant approach is to perform bone marrow transplantation (BMT) after *in vitro* introduction of gRNAs (Fig 7). There is no doubt that these applications have important implications in developmental and stem cell biology.

**Fig 7.**
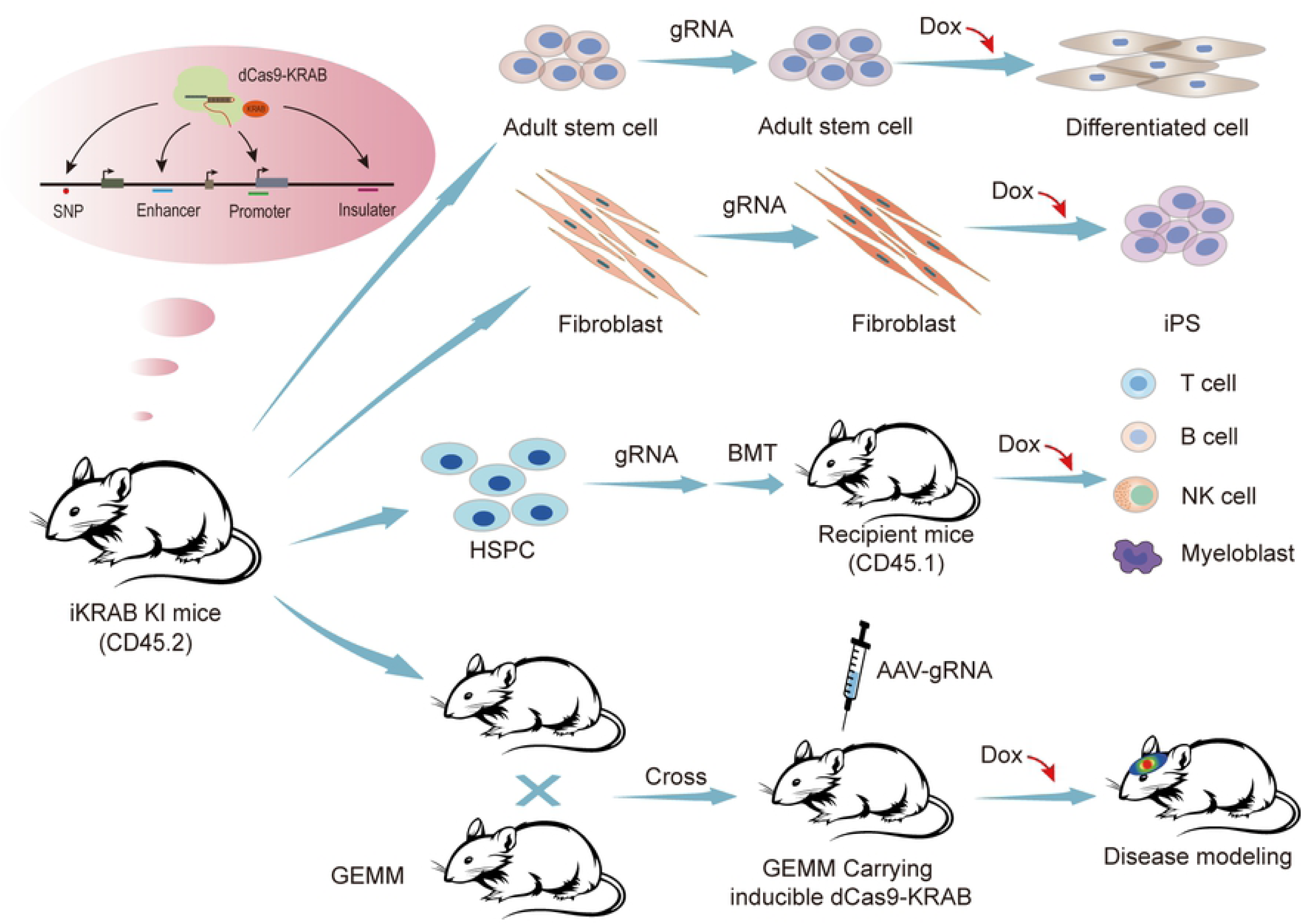
The versatility of the iKRAB KI mouse. Primary cells isolated from the iKRAB KI mice, after being transduced by gRNAs *in vitro*, can be used for following *ex vivo* or *in vivo* functional studies or screens. BMT represents Bone Marrow Transplantation. Crossing the iKRAB KI mouse with other Genetically Engineered Mouse Model (GEMM), we can deliver gRNAs targeting conserved regions harboring human SNPs for complex disease modeling.

More attractively, the iKRAB KI mouse may be valuable for modeling human diseases. Emerging evidences from genome-wide association studies (GWAS) have shown that the vast majority of disease-associated single-nucleotide polymorphisms (SNPs) are located in the non-coding genomic regions. However, the causal-effect relationship between these non-coding mutations and phenotypes or diseases could hardly be established until the development of CRISPR-mediated genome- and epigenome-editing technologies. However genome-editing to generate disease-mimicking cell lines or mouse models that harbor patient-specific non-coding mutations is time-consuming and uneconomical. A trial of epigenome editing in advance to establish the regulatory potential of genomic regions harboring possible causal variants has been suggested [42]. Undoubtedly our iKRAB systems are at least helpful for testing the conserved LOF variants. A simple delivery of gRNAs in the iKRAB KI mice will accelerate the functional exploration in living organism that were previously beyond reach. Though our iKRAB KI mouse does not express *Cre* and the tissue-specific CRISPRi cannot be realized so far, the problem can be solved to pre-cross with specific *Cre* mouse. Furthermore, the crossing with existed Genetically Engineered Mouse Model (GEMM) will facilitate modeling of complex human diseases including cancers (Fig 7). Moreover these animal models will hopefully allow further drug screening and potential preclinical trials. In a word, these versatile iKRAB systems enable a wide range of CRISPRi applications in biological and disease processes.

## MATERIALS AND METHODS

### Cell culture

Mouse ESCs were cultured in GMEM with 15% FBS and 100 U/ml leukemia inhibitory factor (LIF) in gelatin-coated plates. In order to maintain the naïve state, mouse ESCs were cultured in serum-free medium with N2 and B27 supplement, LIF, MEK inhibitor PD0325901 (1 μM), GSK3 inhibitor CHIR99021 (3 μM). To switch to the primed state, ESC medium was switched to serum-free medium with N2 and B27 supplement, plus Fgf2 (12 ng/ml) and activin A (20 ng/ml) as described [25].

### Cloning and plasmid preparation

The fragment of dCas9-KRAB was amplified from the plasmid (Addgene #50917) and introduced into Gateway Entry vector pCR8/GW/TOPO (Invitrogen) following the manufacturer’s protocol and verified by sequencing. The right donor was subcloned into the destination vector p2Lox-FLAG [18] by Gateway technology (Invitrogen).

### gRNA Design and Cloning

To minimize off-targets, gRNAs were designed at the following website http://crispor.tefor.net/. Synthesized oligonucleotides were annealed and cloned into pLX-sgRNA vector following protocol from Addgene. The sequences of all the gRNAs are listed in S2 Table.

### Generation of iKRAB ESC line

After 16 hrs of Dox treatment, A2Loxcre mouse ESCs [17] were transfected with p2Lox- FLAG-dCas9-KRAB plasmids using Lipofectamine 3000 (Thermo), followed by G418 selection (50 μg/mL) for 7 days. Individual colonies were isolated after approximately 10 days, expanded and screened by PCR for inserted sequence. PCR primers are (F: TTACCACTCCCTATCAGTGATAG; R: AGGAAGCTCTCTTCCAGCCTATG). And the inducible expression of FLAG-dCas9-KRAB protein was confirmed by Western Blot assay.

### RT-qPCR

Total RNA was extracted with TRIZOL (Invitrogen). Reverse transcription and quantitative real-time PCR were performed as described [43]. Gene expression was determined relative to *RPLPO* or *Gapdh* using the ΔCt method. The primers are listed in S3 Table.

### ChIP-qPCR

Chromatin preparation was performed as previously described [44]. Briefly, after crosslinking with 1% formaldehyde for 10 min at room temperature and then quenching with 0.125 M glycine for another 5 min, cells were washed with phosphate-buffered saline (PBS) and lysed in SDS Buffer (100 mM NaCl, 50 mM Tris-Cl pH8.1, 5 mM EDTA pH 8.0, 0.5% SDS, protease inhibitors). Nuceli resuspend in appropriate volume of ice-cold IP Buffer (100 mM NaCl, 50 mM Tris-Cl pH8.1, 5 mM EDTA pH 8.0, 0.3% SDS, 1.0% Triton X-100) was sonicated using a BioRuptor sonicator (Diagenode), followed by centrifugation at 16,000×g for 20 min at 4°C. Chromatin was then divided in different aliquots that were incubated overnight at 4°C with primary antibodies. Next, 30 μl protein G magnetic beads were incubated with the reaction for 3 hrs at 4°C. Beads were washed 3 times with high salt buffer (1% Triton X-100, 0.1% SDS, 500 mM NaCl, 2 mM EDTA, pH 8.0, 20 mM Tris-HCl, pH 8.0). After reversal of the crosslinking, ChIP DNA was purified for qPCR analysis. The primers are listed in S4 Table.

### NPC and Neuronal differentiation

The differentiation was performed following a previous protocol [45]. Briefly, ESCs were cultured in differentiation medium (GMEM medium, 15% FBS, non-essential amino acids, β-mercaptoethanol, L-glutamine, penicillin/streptomycin, sodium pyruvate) to form embryonic bodies (EB) by hanging drops. After 4 days, EBs were collected and cultured on bacteriological Petri dishes with 5 μM ATRA for another 2 days. Then EBs were digest and seeded on 0.1% gelatin-coated plates for another 2 days with RA. After 8days NPC stage, cells were cultured in N2 medium (DMEM/F12 medium with 3 mg/ml glucose, 1/100 N2 supplement, 10 ng/ml bFGF, 50 U/ml pen/strep, 1 mM/L-glutamine) for another 4 days without ATRA to generate neurons. The derived neurons were maintained in complete neurobasal medium before proceeding for analysis.

### PDLSC extraction and differentiation

PDLSCs were isolated from 8 weeks iKRAB KI mice. Clipping four incisors and the surrounding gingival tissue of mice. Repeated washing with PBS. Tissues were digested in Collagenase I and dispase for 1 hr and shaked every 15 min in 37°C. The reaction was then stopped with the same amount of serum, followed by centrifuge and cell seeding on culture dish with StemRD MSC medium for 10 days. For PDLSC differentiation, PDLSC were cultured in osteogenic culture medium (10 mM Sodium b-glycerophosphate, 100 μg/L Vitamin C, 10 nM Dexamethasone) for 5 days [32]. Alkaline phosphatase staining was performed according to the protocol from Beyotime Box.

### Lentivirus and AA preparation

All lentiviruses were generated as previously described [43]. Briefly, lentiviral backbone expressing single or multiplex gRNAs with pAX8 (packaging) and pCMV-VSVG (envelope) plasmids were co-transfected into 293FT cells. After 48 hrs, virus supernatants were harvested, filtered and incubated with iKRAB ESCs or primary cells from the iKRAB KI mice. For AAV2 production, HEK293 cells were transfected with the pAAV2 plasmid expressing gRNAs, helper plasmid pDF6, and PEI Max (Polysciences, Inc. 24765-2). At 72 hr posttransfection, the cells were rinsed and pelleted via low-speed centrifugation. Afterward, the viruses were applied to HiTrap heparin columns (GE Biosciences 17-0406-01) and washed with a series of salt solutions with increasing molarities. During the final stages, the eluates from the heparin columns were concentrated using Amicon ultra-15 centrifugal filter units (Millipore).

### Chromosome conformation capture (3C)

3C assays were performed according to previous reports [46]. Cells were crosslinked with 1% formaldehyde for 10 min and quenched with 0.125 M glycine for 5 min. Fixed cells were resuspended in lysis buffer (50 mM Tris–HCl, pH 7.5, 150 mM NaCl, 0.5% NP-40) for 1 hr on ice. Nuclei were resuspended in 0.5 mL of 1X restriction buffer with 0.3% SDS and incubated at 37°C for 1 hr. After that, Triton X-100 was added to a final concentration of 1.8% followed by 20 min incubation at 37°C with shaking. Afterwards, chromatin was digested with HhaI overnight at 37°C. Restriction enzymes was inactivated by adding SDS to a final concentration of 1.6% and incubating the mixture for 20 min at 65°C while shaking. The digested chromatin was diluted 10 times and transferred to new tubes. Then digested chromatin was ligated with 100 U of T4 DNA ligase (with Triton X-100 at a final concentration of 1%) for 8-14 hrs at 16°C. Ligated chromatins were de-crosslinked with 300 mg of Proteinase K and incubated at 65°C overnight, followed by RNase A treatment for 1 hr at 37°C. DNA was then purified by phenol/chloroform extraction followed by ethanol precipitation and re-suspension in water. 3C primers were designed in HhaI fragments located within the enhancer and promoter regions of interest. In addition, we also designed a pair of PCR primers to amplify a ∼200 bp fragment without intervening HhaI sites at the *Sox2* locus, which was used as a loading control. The primers are listed in S5 Table.

### SgRNA library preparation and CRISPRi screening

For each transcript, three to five sgRNAs were designed using CRISPRseek within 500 bp upstream and downstream of the TSS (including alternative TSS). sgRNA sequences that contained BsmbI restriction sites were excluded. The oligonucleotide library was synthesized, annealed, amplified and ligated into the linearized pKLV-U6gRNA-EF(BbsI)- PGKbsd2ABFP vector (modified from addgene #62348 by replacing puro with bsd resistance cassette) for lentivirus packaging. The iKRAB ESCs were transduced with pooled lentiviral sgRNA with an MOI<0.3. After selection of BSD (10 µg/ml) for 4 days, the transduced cells proceeded for NPC differentiation. A803467 (8 nM) and Dox (1 µg/ml) was treated at 4 days after differentiation. 3*10^6^ cells each from the input and survived cells were harvested for genomic DNA extraction 8 days after differentiation and selection. the double sgRNA-encoding regions were then amplified by PCR followed by NGS library preparation (Vazyme cat.TD503-01) and sequencing on an Illumina Hiseq-2500. The amplification primers for the library construction and NGS summarized in S6 Table)

### Generation of animal models and AAV delivery

ESCs was injected to embryonic day 3.5 mouse blastocysts to obtain the founder mice. Chimeric founder mice were bred with C57BL/6 mice, and offsprings with germline transmission were genotyped and intercrossed to generate iKRAB heterozygous or homozygous KI mice. All mouse experiments were performed under protocols approved by Health guidelines of Tianjin Medical University Institutional Animal Use and Care Committee. Animals were fed standard chow diets with access to drinking water ad libitum while housed under a 12-hr light-dark cycle. Six weeks old iKRAB KI mice were injected at multiple sites in the *tibialis anterior* with AAV-Tfam (virus titer: 8.1E12) with 30 µl per mice. Injections were carried out under general anesthesia. After 2 weeks, animals were fed by 5% sucrose water with 1 mg/ml Dox for 1 month. Mice were humanely killed via cervical dislocation and the muscles were rapidly excised.

### IF

Cells were seeded onto slides followed by different treatments and proceeded for IF analysis as previously described [47]. The primary antibodies were listed in S7 Table. The muscle tissue preparation and IF analysis were performed as described [48]. Muscles isolated from tendon to tendon and covered by optimum cutting temperature (OCT) cryoprotectant (Sakura) were rapidly passed in liquid nitro-gen-cooled isopentane (VWR) for 1 min, and left at -80°C until processed. Frozen samples were cryosectioned at 8 μm thickness using a Leica CM1860 cryostat. Then sections were fixed for 10 min with 4% paraformaldehyde in PBS, washed in PBS and blocked in a solution consisting of 1% Tween-20, 5% BSA and PBS for 1 hr. Then the sections were incubated in anti-laminin antibodies overnight at 4 °C. After two washes with PBS-1% Tween-20, samples were incubated with secondary antibodies (1:200, ZSGB-BIO, Alexa-Fluor-594) in PBS for 2 hrs, followed by 5 min incubation in DAPI nuclear stain (Life Technologies). Images were captured using a DP72 fluorescence microscope (Olympus).

### Measurement of Myofiber Cross-Sectional Area

The cross-sectional area of the myofibers was calculated on section images obtained from *Tibialis anterior* muscles using ImageJ.

## Abbreviations

AAV: Adeno-Associated Virus;
ALP: alkaline phosphatase;
ChIP: chromatin immunoprecipitation;
CRISPR: Clustered regularly interspaced short palindromic repeat;
Cas9: (CRISPR-associated) 9;
CRISPR: interference (CRISPRi);
dCas9: deactivated Cas9;
Dox: doxycycline;
EB: embryonic body;
ESC: Embryonic stem cell;
hr: hour;
IF: Immunofluorescence;
KI: knockin;
KRAB: Krüppel-associated box;
LOF: loss-of-function;
MSC: mesenchymal stem cell;
NPC: Neural Progenitor Cell;
PDLSC: Periodontal ligament stem cells;
PE: proximal enhancer;
RT-qPCR: quantitative reverse transcription PCR;
sgRNA: single guide RNA;
SNPs: single-nucleotide polymorphisms;
TSS: transcription start site.

## Supporting information

**S1 Fig. Confirmation of the iKRAB ESC line**. (**A**) Genotyping PCR analysis of proper integration of dCas9-KRAB at the designed locus. (**B**) Western blot analysis showing the inducible expression of FLAG-dCas9-KRAB protein by different concentration of Dox. Gapdh served as a loading control.

**S2 Fig. CRISPRi is insufficient at active genes**. (**A**) Oct4 expression levels are downregulated around 50% accompanied with higher rate of Nanog-positive cells. Immunofluorescence staining of Oct4 and Nanog in SL-cultured iKRAB cells containing Oct4#sgRNA-4 treated with or without Dox. The scale bar represents 50 μm. (**B**) RT-qPCR analysis of stable iKRAB ESCs containing sgRNA against *Bap1* showed less than 50% knockdown efficiency after 2 days of Dox induction. The binding location of each sgRNA is indicated relative to the TSS of *Bap1* locus. Data are represented as the mean ± SD of replicates (n=3). * p<0.05, two-tailed unpaired *t* test.

**S3 Fig. dCas9-KRAB binds at active and inactive chromatin regions comparably**. ChIP-qPCR analysis of dCas9-KRAB around the TSS of *Oct4* and *Fgf5* with Cas9 and FLAG antibodies respectively. Data are represented as the mean ± SD of replicates (n=3).

**S4 Fig. CRISPRi targeting *Oct4-*PE does not downregulate Oct4 expression in 2i- cultured ESCs**. RT-qPCR analysis of Oct4 expression in stable iKRAB ESCs (2i condition) containing sgRNA against *Oct4-*PE. Data are represented as the mean ± SD of replicates (n=3).

**S5 Fig. CRISPRi targeting *Oct4-*PE hinders the epigenetic changes induced by medium switch**. ChIP-qPCR analysis of epigenomic alterations at proximal enhancer of *Oct4* with or without Dox treatment during switch from 2i to SL conditions. Data are represented as the mean ± SD of replicates (n = 3) (***p<0.001** p<0.01*p<0.05; and two-tailed unpaired *t* test).

**S6 Fig. CRISPRi targeting *Sox2-*PE hinders the RA-induced chromatin interaction**. 3C- PCR analysis of *Sox2* proximal enhancer in designed groups. The primers tested the interaction between *Oct4-*TSS and *Sox2-*PE served as a negative control.

**S1 Table. List of sgRNAs with log10FC>1 and p < 0**.**05**.

**S2 Table. SgRNA sequences**.

**S3 Table. RT-qPCR primers**.

**S4 Table. ChIP-qPCR primers**.

**S5 Table. 3C-PCR primers**.

**S6 Table. SgRNA and NGS library construction-PCR primers**.

**S7 Table. Antibodies in this study**.

### Acknowledgments

We are very grateful to M. Kybe for providing A2LoxCre mouse ESC. We thank the animal, FACS and Imaging facilities at TMU for technical support. This work was supported by the National key research and development program (2017YFA0504102), the National Natural Science Foundation of China (31701126, 31900464), and the Natural Science Foundation of Tianjin Municipal Science and Technology Commission (18JCQNJC82300), Open grant from the Chinese Academy of Medical Sciences (157-Zk19-02) and the Talent Excellence Program from Tianjin Medical University and Research Project of Tianjin Education Commission.

